# Body composition and markers of cardiometabolic health in transgender youth compared to cisgender youth: a cross-sectional study

**DOI:** 10.1101/603019

**Authors:** Natalie J Nokoff, Sharon L Scarbro, Kerrie L Moreau, Philip Zeitler, Kristen J Nadeau, Elizabeth Juarez-Colunga, Megan M Kelsey

## Abstract

**Context:** Up to 1.8% of adolescents identify as transgender and many more seek care, yet the impact of gender affirming hormone therapy (GAHT) on cardiometabolic health is unknown.

**Objective:** To determine insulin sensitivity and body composition among transgender females (TF) and males (TM) on estradiol or testosterone, compared to cisgender females (CF) and males (CM).

**Design:** Pilot, cross-sectional study conducted from 2016-2018.

Setting. Academic regional transgender referral center.

**Participants:** Transgender adolescents on either testosterone or estradiol for at least 3 months were recruited. Nineteen TM were matched to 19 CM and 42 CF on pubertal stage and body mass index (BMI). Eleven TF were matched to 23 CF and 13 TF to 24 CM on age and BMI.

**Main Outcome Measure(s):** 1/[fasting insulin] and body composition (dual-energy absorptiometry, DXA).

**Results:** Total body fat was lower in TM than CF (29±7 vs. 33±7%, p=0.002) and higher than CM (28±7 vs. 24±9%, p=0.047). TM had higher lean mass than CF (68±7 vs. 64±7%, p=0.002) and lower than CM (69± vs. 73±8%, p=0.029). Insulin sensitivity was not different between the groups.

TF had lower body fat than CF (31±7 vs. 35±8%, p=0.033) and higher than CM (28±6 vs. 20±10, p=0.001). TF had higher lean mass than CF (66±6 vs. 62±7%, p=0.032) and lower than CM (69±5 vs. 77±9%, p=0.001). TF were more insulin resistant than CM (0.078±0.025 vs. 0.142±0.064, p=0.011).

**Conclusions:** Transgender adolescents on GAHT have significant differences in body composition compared to cisgender controls, with a body composition intermediate between BMI-matched cisgender males and females. These changes in body composition may have consequences for the cardiometabolic health of transgender adolescents.

**Precis:** Transgender youth on gender affirming hormone therapy have differences in their percent fat and lean mass compared to cisgender (non-transgender) youth.

## Introduction

In the United States (U.S.), 0.7%-1.8% of youth identify as transgender (defined as gender identity that is different or opposite from sex at birth) (1,2). Some transgender youth will start gender affirming hormone therapy (GAHT); referrals to centers and providers specializing in this care are rising (3). For patients with a diagnosis of gender dysphoria, gonadotropin-releasing hormone analogues (GnRHa) may be started at Tanner 2 pubertal development and testosterone or estradiol started later in adolescence or adulthood (4). Available data in adults, mostly from Europe, show that transgender women treated with estradiol have a higher incidence of strokes and venous thromboembolism (VTE) than both cisgender women and men (those whose gender identity corresponds with sex at birth) (5). Both transgender women treated with estradiol and transgender men treated with testosterone have a higher incidence of myocardial infarction (MI) than cisgender women (5). Furthermore, there are sex differences in heart disease; cisgender men have a higher prevalence of and death rate from heart disease than cisgender women (6). A meta-analysis of markers of cardiometabolic health and risk in transgender adults on GAHT showed changes in lipid parameters for individuals on hormone therapy, but the data on outcomes such as MI, stroke, VTE and death were sparse (7). Another meta-analysis of longitudinal studies of transgender adults on GAHT showed that both transgender men and women gain weight on GAHT. Specifically, transgender women have an increase in body fat and a decrease in lean mass, whereas transgender men have the opposite (8).

There are many gaps in the literature including: 1) virtually no data on adolescents starting hormone therapy, despite this being a rapidly increasing population with therapy started at younger ages than in the past (4), 2) sparse data from the U.S., where different medications are available and where there may be differences in treatment or approach to care and 3) very few studies compare data to cisgender individuals. The present study addresses these gaps by evaluating markers of cardiometabolic health in transgender adolescents in the U.S. on hormone therapy compared to cisgender adolescents.

The aims of our cross-sectional pilot study are to evaluate insulin sensitivity and body composition among adolescent transgender females (TF) and males (TM) receiving estradiol or testosterone treatment, respectively, matched to cisgender females (CF) and males (CM) of the same body mass index (BMI) and either age or pubertal stage.

## Materials and methods

### Participants

Transgender youth up to age 21 years were recruited between 2016 to 2018 from the TRUE Center for Gender Diversity at Children’s Hospital Colorado. Participants were eligible if they had been on either testosterone or estradiol treatment for at least 3 months. Youth were excluded if they had significant medical or psychiatric comorbidities (including diabetes or antipsychotic treatment). The study was approved by the Colorado Multiple Institutional Review Board and consent and/or assent were obtained from all participants and their guardian (for those under age 18 years).

Twenty-one TMs and 14 TFs participated. The electronic medical records of the transgender participants were reviewed and the start dates for testosterone or estradiol and/or GnRHa treatment and duration of treatment was collected.

Data on healthy cisgender controls were obtained from two studies performed at the same institution: the Colorado RESistance to InSulin in Type 1 ANd Type 2 diabetes (RESISTANT) study and the Health Influences in Puberty (HIP) study. Inclusion criteria for RESISTANT have previously been described (9). In RESISTANT, pubertal adolescents ages 12-19 years were recruited. In the HIP study, adolescents in early puberty (Tanner Stage 2-3) who were either normal weight or obese were recruited between 2009 and 2015 and through local pediatric practices. Presence of diabetes, prediabetes, or medications affecting glucose metabolism was exclusionary. Body composition was measured using dual-energy absorptiometry (DXA) in all studies. Participants in each study had a puberty exam performed by a pediatric endocrinologist.

### Research visit

All transgender participants had a research visit in the morning in the University of Colorado Anschutz Medical Center (UC-AMC) pediatric Clinical Translational Research Center (CTRC) after an overnight fast. For participants taking testosterone, the visit was performed prior to their testosterone injection to obtain a trough value. Pubertal staging was performed by a pediatric endocrinologist using the standards of Tanner and Marshall for breast development (using inspection and palpation), testicular development and pubic hair (10,11); testicular volume was assessed using a Prader orchidometer and assigned a Tanner stage equivalent as follows: Tanner 1 < 4 mL; Tanner 2 ≥ 4 mL and <8mL; Tanner 3 ≥ 8 and < 12 mL; Tanner 4 ≥ 12 and ≤ 15 mL; Tanner 5 >15 mL. Several participants declined genital exam. TM who had already undergone chest masculinizing surgery were assigned Tanner 5 breast development for this analysis. Height was measured on a Harpenden stadiometer and weight on a digital electronic scale. Height and weight were recorded to the nearest 0.1 centimeter and kilogram, respectively. Blood pressure was measured, sitting for at least 5 minutes, with an age-appropriate cuff. Weight, height and blood pressure were each measured twice and averaged. BMI was calculated by weight in kilograms divided by height in meters squared. As all participants were under 20 years, pediatric norms for BMI were used (percentile, where 5 to <85% is normal weight, 85 to <95% is overweight and ≥95% is obese).

Participants filled out a demographic questionnaire and all study data were managed using REDCap electronic data capture tools hosted at the University of Colorado (12).

Fasted blood samples were drawn in the morning and body composition was measured by DXA.

### Laboratory assays

Serum/plasma fasted blood samples were assayed for: glucose, insulin, lipid panel, aspartate aminotransferase (AST), alanine aminotransferase (ALT), hematocrit, hemoglobin A1C, leptin, sex hormone-binding globulin (SHBG), estradiol and total testosterone. Laboratory assays were performed by the UC-AMC CTRC Core Laboratory and the UC Health (UCH) Clinical Laboratory. Glucose was measured by enzymatic UV testing (AU480 Chemistry Analyzer, Beckman Coulter, Brea, CA), inter- and intra-assay coefficient of variation (CV) 1.44% and 0. 67%, respectively and sensitivity 10 mg/dL. Insulin was measured by radioimmunoassay (EMD Millipore, Darmstadt, Germany) with inter- and intra-assay CV 9.8% and 5.2%, respectively, and sensitivity 3 uIU/mL. Leptin was measured by radioimmunoassay (Millipore, Darmstadt, Germany); inter- and intra-assay CV 5.8% and 5.9%, respectively, and sensitivity 0.5 ng/mL.

Testosterone, estradiol, and SHBG were measured by chemiluminescence (Beckman Coulter, Brea, CA). Testosterone inter- and intra-assay CV was 5.1% and 2.1%, respectively, and sensitivity 17 ng/dL; estradiol inter- and intra-assay CV 8.2% and 4.3%, respectively, and sensitivity 10.0 pg/mL; SHBG inter- and intra-assay CV 5.7% and 3.6%, respectively, and sensitivity 3 nmol/L. In HIP and RESISTANT, testosterone and insulin were measured on the same platforms in the same lab.

Total cholesterol, triglycerides (TG) and high-density lipoprotein (HDL) were directly measured and low-density lipoprotein (LDL) was calculated using the Friedewald formula (for units in mg/dL) (13). Insulin sensitivity was estimated by the inverse of the fasting insulin concentration (1/[fasting insulin]), which is correlated with insulin sensitivity measured with a hyperinsulinemic euglycemic clamp (14); lower values for 1/insulin indicate worse insulin sensitivity. Free androgen index (FAI) was calculated as the ratio of total testosterone to SHBG ([testosterone/SHBG]*100) (15).

### Statistical analysis

TM participants (n=21, ages 15.1-19.8 years) were matched on body mass index (BMI) percentile to cisgender controls, allowing for a +/− 6% difference using the GREEDY matching algorithm (16). All controls and all but 2 TM were Tanner Stage 5 (missing data for 2 TM participants). TM participants (n=19) were matched to CM (n=19, ages 13.1-19.7 years) and CF (n=42, ages 11.7-18.9 years) with a range of 1 to 3 CF matches per TM case (1:1 n=19, 1:2 n=14, 1:3 n=9). The 19 TM participants to the CM and CF cases were not the same 19 based on the available matches.

TF (n=14, ages 14.5-19.4 years) were matched (2 were unable to be matched due to lack of matching weight controls) to cisgender youth on age (within a year) and BMI in two phases. The first phase performed a one-to-one match using the GREEDY algorithm with BMI percentile +/− 12.5% and age within a year. The second phase identified additional matches with BMI percentile in the same category and age within a year. TF (n=13) were matched to CM (n=24, ages 14.5-19.8 years) with a range of 1 to 5 matches per TF case (1:1 n=8, 1:2 n=2, 1:3 n=1, 1:4 n=1, 1:5 n=1). TF (n=11) were matched to 23 CF (ages 14.2-18.3) with a range of 1 to 4 matches per TF case (1:1 n=5, 1:2 n=2, 1:3 n=2, 1:4 n=2). One participant did not have estradiol results.

BMI percentile was defined using CDC criteria: 5 to <85% being normal weight, 85 to <95% as overweight and >95% as obese. The 2000 CDC Growth Charts were used to calculate percentiles (17).

Because the AST and ALT for the cisgender participants were measured by a different assay, a correction factor was applied to the transgender participants: corrected AST = (measured AST +14.374)/0.8334 and corrected ALT = (measured ALT +10.058)/0.9319. Deming regression was used to build a regression model and determine the correction needed to make the results equivalent using the parameter estimates obtained from the regression.

Tests of differences between TF and controls and TM and controls were performed by running a mixed linear regression model with a random effect for the matched set. Compound symmetry was used for the covariance structure and the restricted maximum likelihood method was used to estimate the covariance parameters. Comparisons with TM were adjusted for age in years because we did not match on age for the TM group. Outcomes that were not normally distributed were log transformed and the log-transformed p-values are presented. For group comparisons in which there was more than one cisgender match for the transgender participant, means of the matched set are presented.

Our preferred approach would have been to match on pubertal stage and adjust for age for all groups, given that insulin resistance changes with pubertal stage (18). Because all TM participants were Tanner 5, we adjusted for age. However, for the TF cohort, more participants had received a GnRHa and had a lower Tanner stage than similar aged controls, so we were unable to match on Tanner stage. Therefore, we matched on age, to try and account for differences in insulin sensitivity throughout puberty. We did not adjust for pubertal stage because of missing data.

Spearman correlations were performed to evaluate for correlations between: 1) % body fat and leptin and 2) % lean mass and inverse of fasting insulin. Analyses were conducted using SAS version 9.4 (SAS Institute, Cary, NC). P-values <0.05 were considered significant. We did not correct for multiple comparisons since this was a pilot study and all findings are considered exploratory.

## Results

Demographics of the overall group are presented in Tables 1 and 2. TM were on an average testosterone dose of 217 ± 88 mg/month for an average treatment duration of 11.2 ± 5.9 months. Twelve (57%) were using intramuscular (IM) injections and 9 (43%) were using subcutaneous (SQ) injections. None were on a GnRHa at the time of the study visit but one participant had recently discontinued the GnRHa (length of GnRHa therapy 17.1 months) and therefore may still have been experiencing the effects of the medication. An additional 3 participants had used a GnRHa in the past. Four participants were using a progestin at the time of the study visit (3 on medroxyprogesterone, one with an etonogestrel implant). Six participants had undergone chest masculinizing surgery. None had received any other types of surgeries. TF were taking an average estradiol dose of 1.5 ± 1.0 mg/day with an average treatment duration of 12.3 ± 9.9 months (five on oral, 9 on sublingual). Four were on a GnRHa at the time of the study visit and a total of 6 had been on a GnRHa in the past.

**Table 1:** Demographics of transgender males and cisgender females and males.

|  | Transgender male<br>(n=21) | Cisgender female<br>(n=42) | Cisgender male<br>(n=19) |
| --- | --- | --- | --- |
| Age (years) | 17.0 ± 1.4 | 15.2 ± 1.9 | 15.3 ± 1.6 |
| Race |  |  |  |
| White | 15 (71) | 28 (67) | 14 (74) |
| Asian | 1 (5) | 3 (7) | 2 (11) |
| African-American | 0 | 9 (21) | 2 (11) |
| Native American/ Alaska Native | 1 (5) | 1 (2) | 0 |
| More than one race | 3 (14) | 0 | 1 (5) |
| Unknown/not reported | 1 (5) | 1 (2) | 0 |
| Ethnicity |  |  |  |
| Hispanic | 7 (33) | 14 (33) | 3 (16) |
| Not Hispanic | 14 (67) | 26 (62) | 16 (84) |
| Unknown/not reported | 0 | 2 (5) | 0 |
| BMI category |  |  |  |
| Normal weight | 14 (67) | 25 (60) | 14 (74) |
| Overweight | 5 (24) | 7 (17) | 1 (5) |
| Obese | 2 (10) | 10 (24) | 4 (21) |
| Pubic hair Tanner stage |  |  |  |
| 1 | 0 | 0 | 0 |
| 2 | 0 | 0 | 0 |
| 3 | 0 | 0 | 1 (5) |
| 4 | 1 (5) | 8 (19) | 7 (37) |
| 5 | 14 (67) | 6 (14) | 9 (47) |
| Missing | 6 (29) | 28 (67) | 2 (11) |
| Breast/testicular Tanner stage |  |  |  |
| Stage 5 | 19 (90) | 42 (0) | 19 (100) |
| Missing | 2 (10) | 0 | 0 |
| Had menarche | 20 (95) | 26 (62) | --- |
| Currently having menses | 2 (10) |  | --- |
| Age of menarche | 11.9 ± 1.1 | 12.4 ± 1.4 | --- |
| Family history* |  |  |  |
| Hypertension | 13 (68) | 12 (86) | 15 (88) |
| Hypercholesterolemia | 10 (52) | 11 (79) | 14 (82) |
| Type 2 diabetes | 10 (52) | 11 (46) | 8 (47) |

Values above represent the entire cohort used and are either presented as mean ± standard deviation or n (%).
Different TM participants were used to match to different cisgender males (CM) or females (CF) based on the ideal match for BMI. Mean ± SD or n (%). \*For CM and CF, percentages are given out of total number of reported values (non-missing). Family history data was missing for several conditions. For CF, there was missing family history of hypertension and hypercholesterolemia for 28 and type 2 diabetes for 18 participants. For CM, there was missing family history of hypertension, hypercholesterolemia and type 2 diabetes for 2 participants.

**Table 2:**
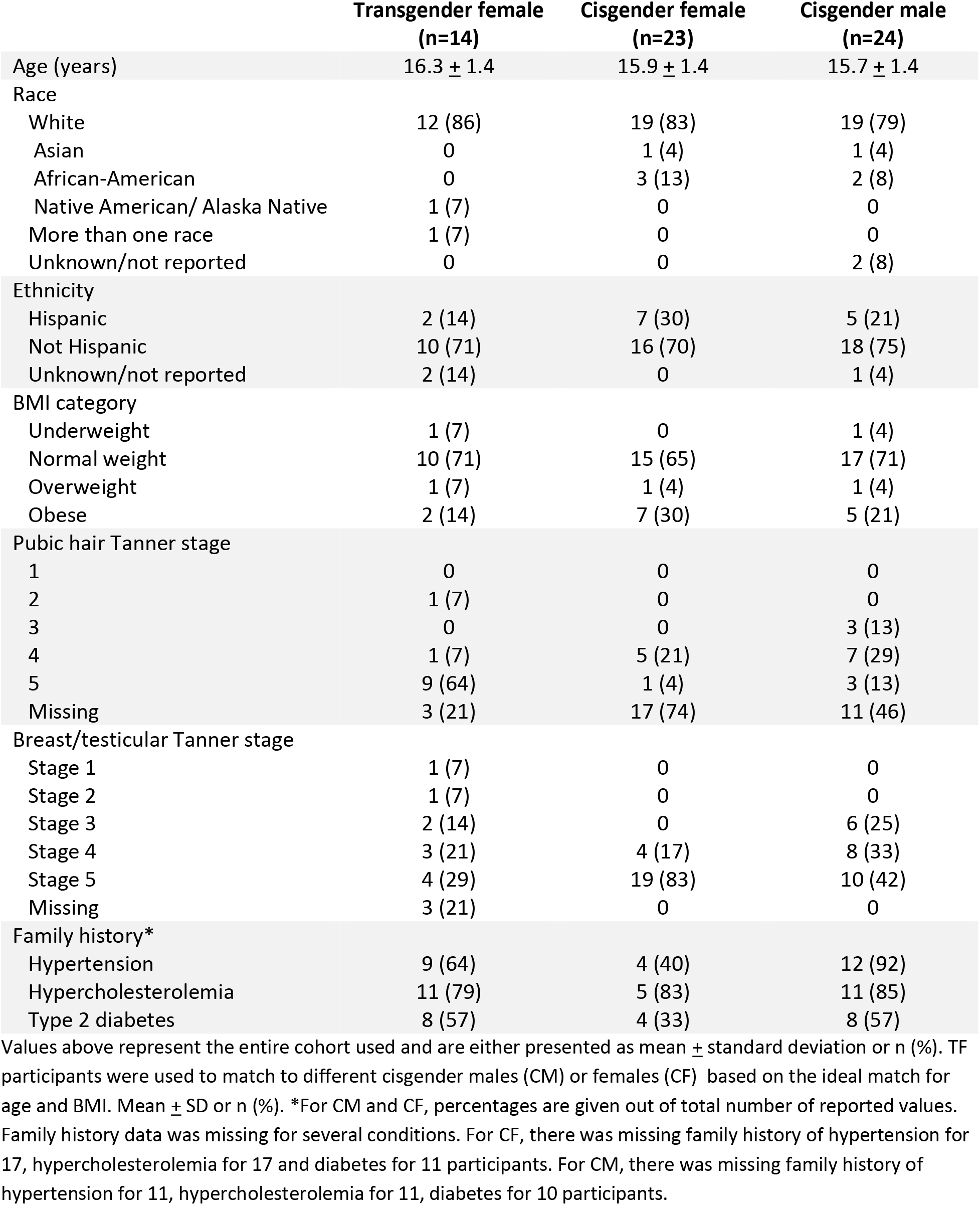
Demographics of transgender females and cisgender females and males.

### Transgender males compared to cisgender females

Markers of cardiometabolic health and hormone concentrations for TM are displayed in Table 3 and Figure 1. TM had a higher AST (p=0.001), lower HDL (p=0.043) and lower leptin (p=0.018) than CF. Body composition was significantly different between groups (Table 3, Figure 1a and b). TM had a lower % body fat (p=0.002) and fat mass (p=0.029), and higher % lean tissue (p=0.002) and lean mass (p=0.039) than CF. Compared to CF, TM had higher serum testosterone (p<0.001) and FAI (p<0.001), and lower SHBG (p<0.001). There were no differences in insulin sensitivity between TM and CF.

**Figure 1:**
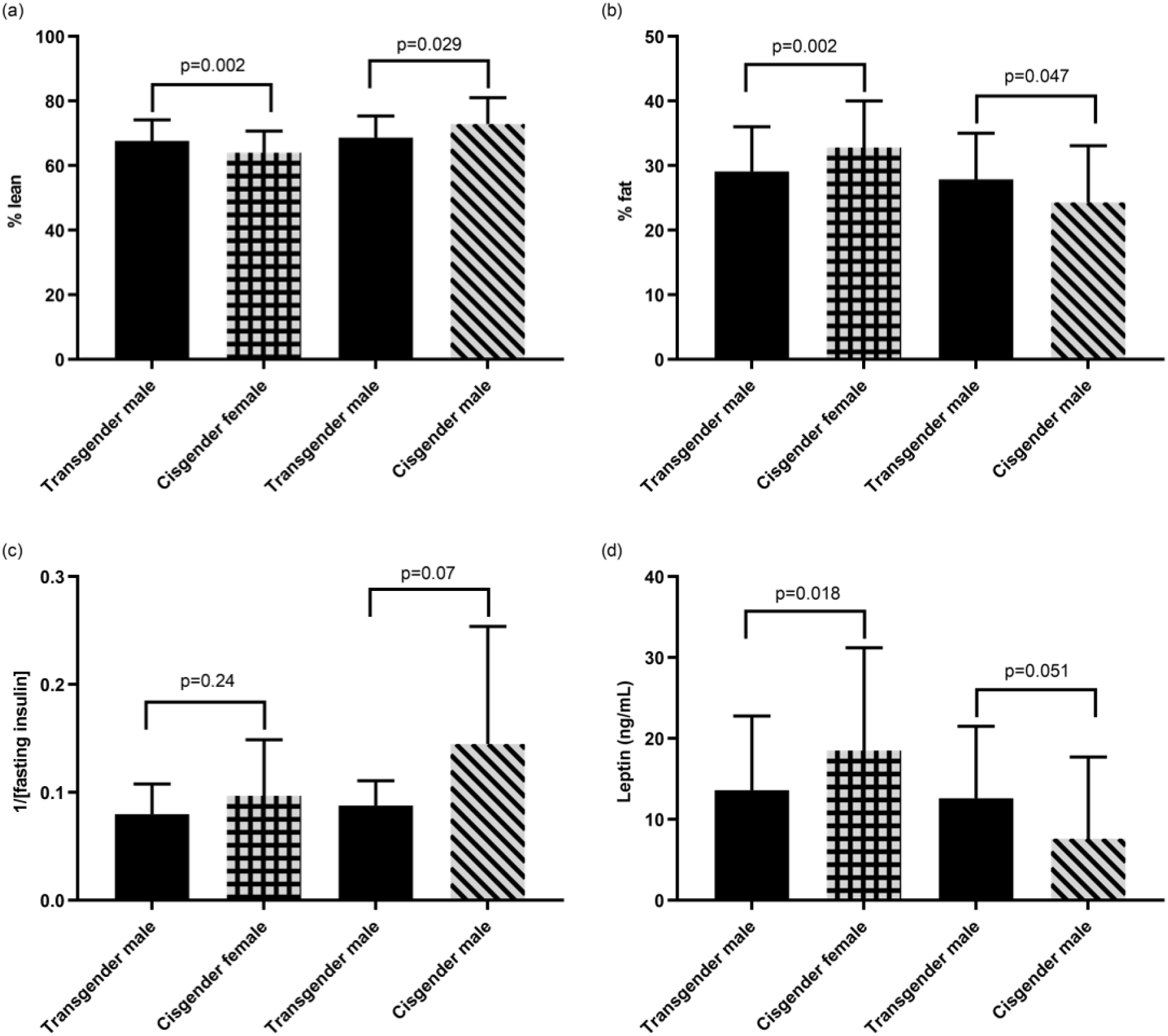
Body composition, insulin sensitivity and leptin in transgender males and cisgender males and females. Means and standard deviations are presented. Transgender males are presented twice because the same individuals are not compared to both cisgender males and females

**Table 3:** Markers of cardiometabolic health and hormone concentrations for transgender males compared to cisgender males and females.

|  | Transgender male<br>(n=19) | Cisgender female<br>(n=19†) | Transgender male<br>(n=19) | Cisgender male<br>(n=19) |
| --- | --- | --- | --- | --- |
| Age | 16.9 ± 1.4 | 14.9 ± 1.7 | 17.1 ± 1.4 | 15.3 ± 1.6 |
| BMI (%) | 71 ± 22 | 71 ± 21 | 63 ± 28 | 64 ± 28 |
| Systolic BP (mmHg) | 108 ± 9 | 111 ± 8 | 108 ± 9 | 115 ± 13** |
| Diastolic BP (mmHg) | 70 ± 7 | 66 ± 6 | 69 ± 8 | 67 ± 10 |
| Inverse of fasting insulin | 0.080 ± 0.028 | 0.097 ± 0.052 | 0.088 ± 0.023 | 0.145 ± 0.109 |
| Fasting glucose (mg/dL) | 89 ± 5 | 85 ± 6 | 88 ± 5 | 86 ± 10 |
| Hemoglobin A1C (%) | 5.3 ± 0.2 | 5.2 ± 0.2 | 5.3 ± 0.3 | 5.3 ± 0.3 |
| AST (U/L) | 39 ± 5 | 29 ± 8** | 39 ± 4 | 36 ± 16 |
| ALT (U/L) | 26 ± 5 | 26 ± 7 | 25 ± 6 | 34 ± 17*** |
| Total cholesterol (mg/dL) | 147 ± 16 | 153 ± 29 | 143 ± 19 | 146 ± 22 |
| Triglycerides (mg/dL) | 75 ± 21 | 100 ± 45 | 76 ± 23 | 91 ± 30 |
| HDL (mg/dL) | 40 ± 5 | 46 ± 7* | 41 ± 5 | 46 ± 9 |
| LDL (mg/dL) | 92 ± 16 | 87 ± 22 | 87 ± 19 | 82 ± 19 |
| Total estradiol (pg/mL) | 43 ± 23 | 63 ± 40 | 46 ± 22 | 24 ± 11** |
| Total testosterone (ng/dL) | 363 ± 220 | 39 ± 13*** | 378 ± 219 | 445 ± 152 |
| SHBG (nmol/L) | 24 ± 11 | 47 ± 25*** | 26 ± 11 | 36 ± 13 |
| Free androgen index | 65 ± 47 | 4 ± 2*** | 64 ± 47 | 48 ± 16 |

Values are given as mean ± SD. Transgender males are presented twice because the same 19 individuals are not compared to both cisgender males and females. BP= blood pressure, †means of the matched set, \*p<0.05, \*\*p≤0.01, \*\*\*p≤0.001 (p-values represent significance from log-transformed variables when relevant)

### Transgender males compared to cisgender males

Compared to CM, TM had a lower systolic blood pressure (p=0.005) and ALT (p<0.001). Body composition was significantly different between groups (Table 3, Figure 1a and 1b). TM had a higher % body fat (p=0.047), and lower % lean tissue (p=0.029) and lean mass (p<0.001) than CM. Compared to CM, TM had higher serum estradiol (p=0.004). There were no differences in insulin sensitivity between TM and CM.

### Transgender females compared to cisgender females

Markers of cardiometabolic health and hormone concentrations for TF are shown in Table 4 and Figure 2. TF had a higher AST than CF (p<0.001). Body composition was significantly different between groups (Table 4, Figure 2a and 2b). Compared to CF, TF had lower % body fat (p=0.033), and higher % lean tissue (p=0.032) lean mass (p=0.004), total testosterone (p<0.001) and FAI (p=0.002) than CF. There were no differences in insulin sensitivity between TF and CF.

**Figure 2:**
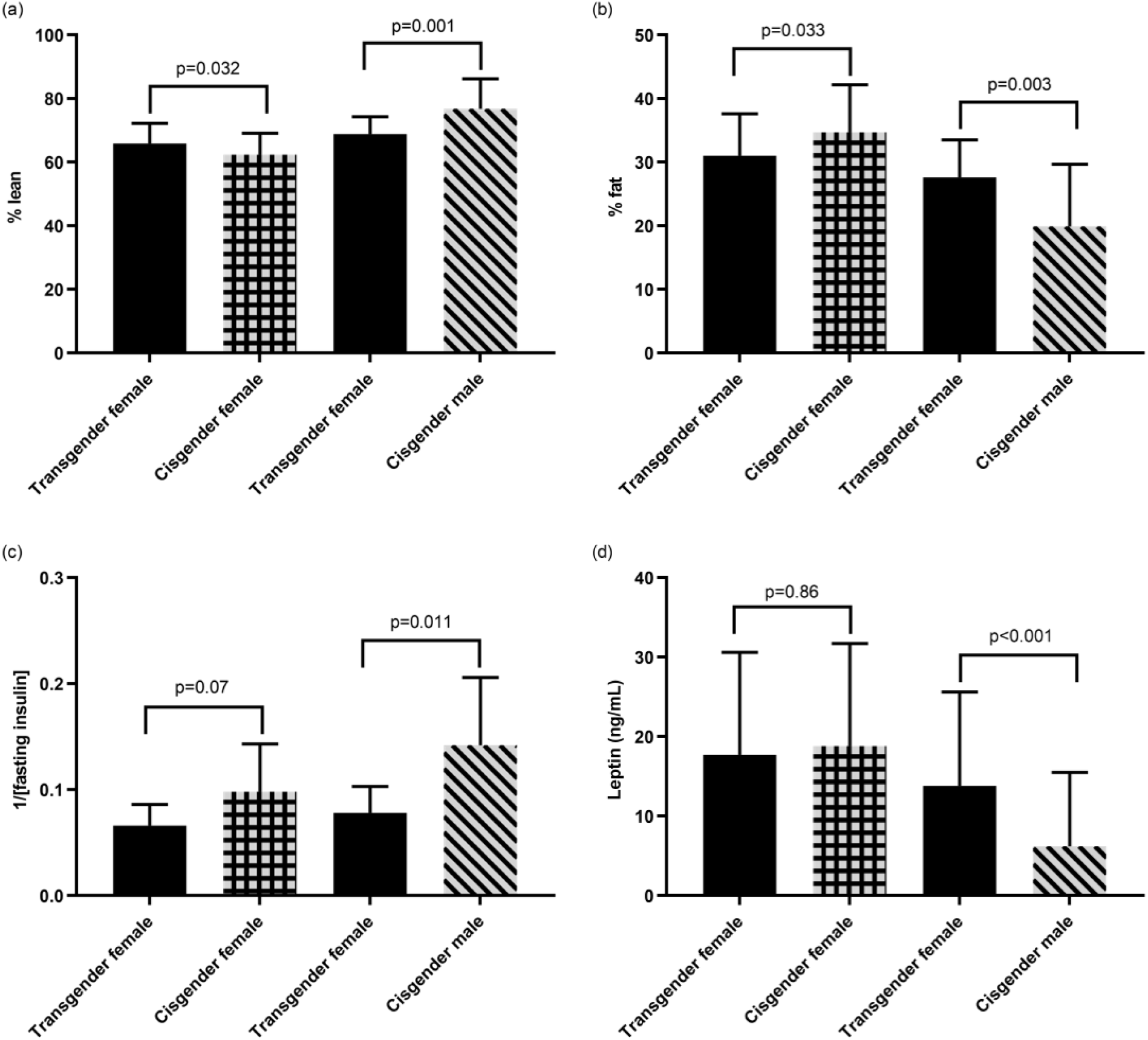
Body composition, insulin sensitivity and leptin in transgender females and cisgender males and females. Means and standard deviations are presented. Transgender females are presented twice because the same individuals are not compared to both cisgender males and females

**Table 4:**
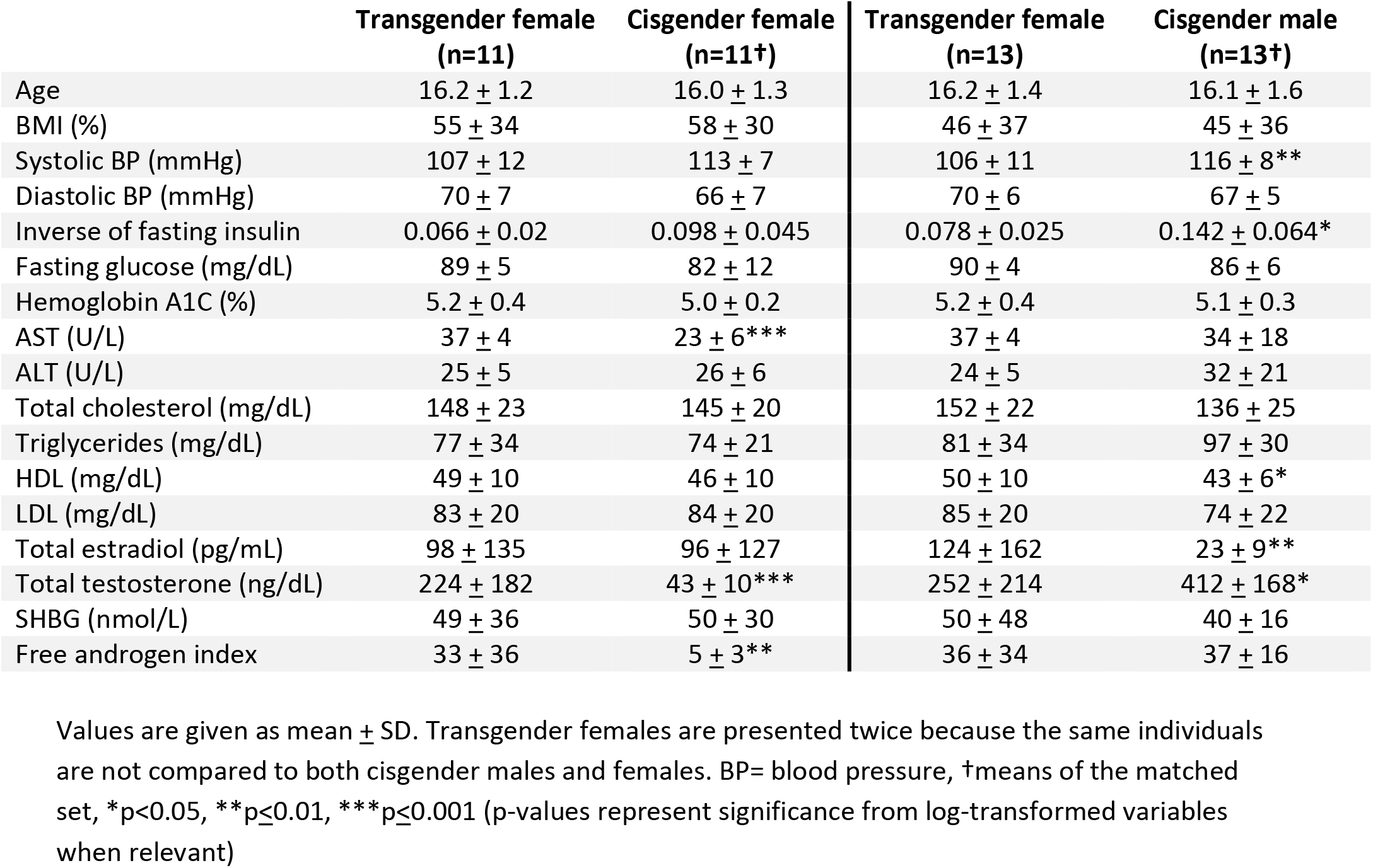
Markers of cardiometabolic health and hormone concentrations transgender females compared to cisgender females and males.

|  | Transgender female<br>(n=11) | Cisgender female<br>(n=11†) | Transgender female<br>(n=13) | Cisgender male<br>(n=13†) |
| --- | --- | --- | --- | --- |
| Age | 16.2 ± 1.2 | 16.0 ± 1.3 | 16.2 ± 1.4 | 16.1 ± 1.6 |
| BMI (%) | 55 ± 34 | 58 ± 30 | 46 ± 37 | 45 ± 36 |
| Systolic BP (mmHg) | 107 ± 12 | 113 ± 7 | 106 ± 11 | 116 ± 8** |
| Diastolic BP (mmHg) | 70 ± 7 | 66 ± 7 | 70 ± 6 | 67 ± 5 |
| Inverse of fasting insulin | 0.066 ± 0.02 | 0.098 ± 0.045 | 0.078 ± 0.025 | 0.142 ± 0.064* |
| Fasting glucose (mg/dL) | 89 ± 5 | 82 ± 12 | 90 ± 4 | 86 ± 6 |
| Hemoglobin A1C (%) | 5.2 ± 0.4 | 5.0 ± 0.2 | 5.2 ± 0.4 | 5.1 ± 0.3 |
| AST (U/L) | 37 ± 4 | 23 ± 6*** | 37 ± 4 | 34 ± 18 |
| ALT (U/L) | 25 ± 5 | 26 ± 6 | 24 ± 5 | 32 ± 21 |
| Total cholesterol (mg/dL) | 148 ± 23 | 145 ± 20 | 152 ± 22 | 136 ± 25 |
| Triglycerides (mg/dL) | 77 ± 34 | 74 ± 21 | 81 ± 34 | 97 ± 30 |
| HDL (mg/dL) | 49 ± 10 | 46 ± 10 | 50 ± 10 | 43 ± 6* |
| LDL (mg/dL) | 83 ± 20 | 84 ± 20 | 85 ± 20 | 74 ± 22 |
| Total estradiol (pg/mL) | 98 ± 135 | 96 ± 127 | 124 ± 162 | 23 ± 9** |
| Total testosterone (ng/dL) | 224 ± 182 | 43 ± 10*** | 252 ± 214 | 412 ± 168* |
| SHBG (nmol/L) | 49 ± 36 | 50 ± 30 | 50 ± 48 | 40 ± 16 |
| Free androgen index | 33 ± 36 | 5 ± 3** | 36 ± 34 | 37 ± 16 |

Values are given as mean ± SD. Transgender females are presented twice because the same individuals are not compared to both cisgender males and females. BP= blood pressure, †means of the matched set, \*p<0.05, \*\*p≤0.01, \*\*\*p≤0.001 (p-values represent significance from log-transformed variables when relevant)

### Transgender females compared to cisgender males

TF were more insulin resistant than CM, with a lower inverse of fasting insulin (p=0.011) and a higher HOMA-IR (p=0.012). TF had a lower systolic blood pressure (p=0.007), and higher HDL (p=0.023) and leptin (p<0.001) than CM. Body composition was significantly different between groups (Table 4, Figure 2a and 2b). Compared to CM, TF had higher % body fat (p=0.003) and fat mass (p=0.004), and lower % lean tissue (p=0.001). TF had higher estradiol (p=0.005) and lower total testosterone (p=0.012) than CM.

### Correlations and other observations

In the pooled population of TM and cisgender controls, the inverse of fasting insulin correlated with % lean mass (r=0.40 [95% CI: 0.19, 0.57], p=0.0004) as well as for TF and controls (r=0.65 [95% CI: 0.46, 0.79], p<0.0001). As expected, leptin correlated strongly with % fat mass for both TM and their controls (r=0.90 [95% CI: 0.84, 0.94], p<0.0001) and TF and their controls (r=0.94 [95% CI: 0.89, 0.97], p<0.0001).

The % lean and % fat mass were remarkably similar for the TF and TM; these groups were not matched on age or BMI or compared to one another.

## Discussion

The present study demonstrates novel observations with regards to cardiometabolic health parameters in transgender youth treated with GAHT. First, both TM and TF had a body composition (defined by percent fat and lean on DXA) that is intermediate between CF and CM. Second, there were both favorable and unfavorable changes in markers of cardiometabolic health for TF and TM compared to BMI-matched cisgender youth. The only group that had a difference in insulin sensitivity were the TF compared to CM, with TF being less insulin sensitive than CM.

Most participants in this study had not received a GnRHa and went through most or all their endogenous puberty before starting testosterone or estradiol. Body composition in the TM may be explained by a female pattern of pubertal fat accrual (19), followed by gains in lean mass with testosterone. Differences in body composition in the TF may be explained by a male pattern of lean mass accrual during puberty, followed by a gain in percent fat and concomitant rise in leptin due to estradiol treatment.

The body composition findings are similar to those observed in adults. In a meta-analysis of longitudinal studies of transgender adults treated with GAHT from 3 to 24 months, all groups had an increase in body weight. TF had an increase in body fat and decrease in lean body mass, whereas TM had the opposite changes (8). Participants were on a variety of hormone regimens, some not routinely used in the U.S., and most individuals had not received a GnRHa. However, in a multi-center retrospective study of adolescents and young adults, only TM had an increase in BMI on testosterone, whereas there were no changes in BMI for TF after starting estradiol (20). In our cross-sectional study, TM and TF had an intermediate body composition, with percent fat and lean mass between CM and CF values. Similarly, leptin, a hormone secreted from fat cells, was higher in TF compared to CM and lower in TM than CF.

Some studies that have examined insulin sensitivity for adults on GAHT have shown decreased insulin sensitivity for both TF and TM and an increase in fasting insulin for TF (21). Another study showed that both TM and TF on GAHT were more insulin-resistant after 4 months of hormone therapy compared to baseline, measured by hyperinsulinemic euglycemic clamp, the gold standard for measuring insulin sensitivity (22). However, TF in that study were treated with ethinyl estradiol (rather than 17β-estradiol, as used in the current study), which is known to have an adverse effect on glucose and insulin (23). Our study was a cross-sectional comparison with control populations, rather than an intra-individual comparison before and after starting treatment, and the TFs were more insulin resistant than CMs but not CFs. In Belgium, both TM and TF have a higher prevalence of type 2 diabetes than CM or CF (24), although the same has not been demonstrated in the U.S. (25).

Many of the studies on cardiometabolic health in transgender individuals involve small cohorts, followed longitudinally. However, it is also important to compare outcomes to appropriately matched cisgender participants, as there are known sex differences in cardiometabolic health that begin to emerge in puberty. Youth become more insulin resistant in puberty (beginning at Tanner 2, peaking at Tanner 3 and returning to near prepubertal values sometime after puberty is completed) (18). There are also sex differences in pubertal insulin resistance, with cisgender girls being more insulin-resistant than boys and, while insulin resistance is strongly related to BMI and body fat, these do not entirely account for the sex differences seen (18). Furthermore, type 2 diabetes in youth is more common among cisgender girls than boys (26,27), a sex difference not seen in adults (28). In adults, worse insulin resistance is associated with incident coronary heart disease, which has been shown to be more common among transgender than cisgender adults (5,29). The impact of GnRHa, testosterone and estradiol started during puberty on present insulin resistance and future risk of type 2 diabetes and heart disease warrants further study.

In TM adults, testosterone therapy is associated with increased LDL and triglycerides and decreased HDL (7). A retrospective study in adolescents and young adults also found that TM on testosterone had a decrease in HDL (20). We found that TM adolescents had lower HDL than CF but no other differences in lipids and no differences compared to CM. TF adults on estradiol had an increase in triglycerides (7). We did not find any statistically significant differences in triglycerides between TF and CM or CF, but TF did have a higher HDL than CM, which is an expected effect of estradiol (30).

As many more transgender youth seek GAHT, it is important to have a better understanding of the impact, not only of testosterone and estradiol, but also of GnRH analogues, on short- and long-term cardiometabolic health. The current study, while cross-sectional, has many strengths. There have been very few rigorously performed studies in transgender youth. One published study, although multi-center, was retrospective, with labs performed at different locations and not necessarily fasting (20). Additionally, although body composition has been measured in transgender adults, this has not, to our knowledge, been done in transgender adolescents. And while most studies in transgender adults have been longitudinal, very few have employed a comparison group with similar characteristics and BMI. There were also several limitations to our study. It was a cross-sectional study, rather than longitudinal, so we do not know about changes before and after GAHT. Some participant data had to be excluded because there were no available matches. The sample size is small and some participants were on a GnRHa and others were not (and the numbers were too small to identify differences between these two groups). Testosterone was not measured by mass spectroscopy in any of the studies.

In conclusion, we show that among transgender adolescents using GAHT for approximately one year, there are significant differences in body composition between transgender and cisgender adolescents, with transgender adolescents having a body composition intermediate between cisgender adolescents of the same BMI. There were also differences in markers of cardiometabolic health between transgender and cisgender youth, the most notable being that TF participants were more insulin-resistant than CM. Based on the results of this pilot study, further exploration is needed to understand the impact of starting testosterone or estradiol treatment in adolescence, with or without prior pubertal blockade, on short- and long-term cardiometabolic health.

## Acknowledgements

The authors wish to thank the participants in the study.

## References

1. Herman JL, Flores AR, Brown TNT, Wilson BDM, Conron KJ. Age of individuals who identify as transgender in the United States. The Williams Institute 2017. Available at: https://williamsinstitute.law.ucla.edu/wp-content/uploads/TransAgeReport.pdf.

2. Johns MM, Lowry R, Andrzejewski J, Barrios LC, Demissie Z, McManus T, Rasberry CN, Robin L, Underwood JM. Transgender Identity and Experiences of Violence Victimization, Substance Use, Suicide Risk, and Sexual Risk Behaviors Among High School Students - 19 States and Large Urban School Districts, 2017. MMWR Morb. Mortal. Wkly. Rep. 2019;68(3):67–71.

3. Chen M, Fuqua J, Eugster EA. Characteristics of Referrals for Gender Dysphoria Over a 13-Year Period. J. Adolesc. Health 2016;58(3):369–371.

4. Hembree WC, Cohen-Kettenis PT, Gooren L, Hannema SE, Meyer WJ, Murad MH, Rosenthal SM, Safer JD, Tangpricha V, T’Sjoen GG. Endocrine Treatment of Gender-Dysphoric/Gender-Incongruent Persons: An Endocrine Society* Clinical Practice Guideline. J. Clin. Endocrinol. Metab. 2017. doi:10.1210/jc.2017-01658.

5. Nota NM, Wiepjes CM, de Blok CJM, Gooren LJG, Kreukels BPC, den Heijer M. The Occurrence of Acute Cardiovascular Events in Transgender Individuals Receiving Hormone Therapy: Results from a Large Cohort Study. Circulation 2019. doi:10.1161/CIRCULATIONAHA.118.038584.

6. National Center for Health Statistics (US). Health, United States, 2017: With Special Feature on Mortality. Hyattsville (MD): National Center for Health Statistics (US); 2019.

7. Maraka S, Singh Ospina N, Rodriguez-Gutierrez R, Davidge-Pitts CJ, Nippoldt TB, Prokop LJ, Murad MH. Sex steroids and cardiovascular outcomes in transgender individuals: a systematic review and meta-analysis. J. Clin. Endocrinol. Metab. 2017. doi:10.1210/jc.2017-01643.

8. Klaver M, Dekker M, Mutsert R, Twisk J, Heijer M. Cross-sex hormone therapy in transgender persons affects total body weight, body fat and lean body mass: a metaanalysis. Andrologia 2017;49(5). Available at: http://onlinelibrary.wiley.com/doi/10.1111/and.12660/full.

9. Bjornstad P, Truong U, Pyle L, Dorosz JL, Cree-Green M, Baumgartner A, Coe G, Regensteiner JG, Reusch JEB, Nadeau KJ. Youth with type 1 diabetes have worse strain and less pronounced sex differences in early echocardiographic markers of diabetic cardiomyopathy compared to their normoglycemic peers: A RESistance to InSulin in Type 1 ANd Type 2 diabetes (RESISTANT) Study. J. Diabetes Complications 2016;30(6):1103–1110.

10. Marshall WA, Tanner JM. Variations in the pattern of pubertal changes in boys. Arch. Dis. Child. 1970;45(239):13–23.

11. Marshall WA, Tanner JM. Variations in pattern of pubertal changes in girls. Arch. Dis. Child. 1969;44(235):291–303.

12. Harris PA, Taylor R, Thielke R, Payne J, Gonzalez N, Conde JG. Research electronic data capture (REDCap)—a metadata-driven methodology and workflow process for providing translational research informatics support. J. Biomed. Inform. 2009;42:377–381.

13. Friedewald WT, Levy RI, Fredrickson DS. Estimation of the concentration of low-density lipoprotein cholesterol in plasma, without use of the preparative ultracentrifuge. Clin. Chem. 1972;18(6):499–502.

14. George L, Bacha F, Lee S, Tfayli H, Andreatta E, Arslanian S. Surrogate estimates of insulin sensitivity in obese youth along the spectrum of glucose tolerance from normal to prediabetes to diabetes. J. Clin. Endocrinol. Metab. 2011;96(7):2136–2145.

15. Al Kindi MK, Al Essry FS, Al Essry FS, Mula-Abed W-AS. Validity of serum testosterone, free androgen index, and calculated free testosterone in women with suspected hyperandrogenism. Oman Med. J. 2012;27(6):471–474.

16. Cormen TH, Leiserson CE, Rivest RL, Stein C. Introduction to Algorithms. MIT Press; 2009.

17. Kuczmarski RJ, Ogden CL, Guo SS, Grummer-Strawn LM, Flegal KM, Mei Z, Wei R, Curtin LR, Roche AF, Johnson CL. 2000 CDC Growth Charts for the United States: methods and development. Vital Health Stat. 11 2002;(246):1–190.

18. Moran A, Jacobs DR Jr, Steinberger J, Hong CP, Prineas R, Luepker R, Sinaiko AR. Insulin resistance during puberty: results from clamp studies in 357 children. Diabetes 1999;48(10):2039–2044.

19. McCarthy HD, Cole TJ, Fry T, Jebb SA, Prentice AM. Body fat reference curves for children. Int. J. Obes. 2006;30(4):598–602.

20. Jarin J, Pine-Twaddell E, Trotman G, Stevens J, Conard LA, Tefera E, Gomez-Lobo V. Cross-Sex Hormones and Metabolic Parameters in Adolescents With Gender Dysphoria. Pediatrics 2017;139(5). doi:10.1542/peds.2016-3173.

21. Gooren LJ, Giltay EJ, Bunck MC. Long-term treatment of transsexuals with cross-sex hormones: extensive personal experience. J. Clin. Endocrinol. Metab. 2008;93:19–25.

22. Polderman KH, Gooren LJ, Asscheman H, Bakker A, Heine RJ. Induction of insulin resistance by androgens and estrogens. J. Clin. Endocrinol. Metab. 1994;79(1):265–271.

23. Sitruk-Ware R, Nath A. Characteristics and metabolic effects of estrogen and progestins contained in oral contraceptive pills. Best Pract. Res. Clin. Endocrinol. Metab. 2013;27(1):13–24.

24. Wierckx K, Elaut E, Declercq E, Heylens G, De Cuypere G, Taes Y, Kaufman JM, T’Sjoen G. Prevalence of cardiovascular disease and cancer during cross-sex hormone therapy in a large cohort of trans persons: a case-control study. Eur. J. Endocrinol. 2013;169(4):471–478.

25. Nokoff NJ, Scarbro S, Juarez-Colunga E, Moreau KL, Kempe A. Health and Cardiometabolic Disease in Transgender Adults in the United States: Behavioral Risk Factor Surveillance System 2015. J Endocr Soc 2018;2(4):349–360.

26. Copeland KC, Zeitler P, Geffner M, Guandalini C, Higgins J, Hirst K, Kaufman FR, Linder B, Marcovina S, McGuigan P, Pyle L, Tamborlane W, Willi S, TODAY Study Group. Characteristics of adolescents and youth with recent-onset type 2 diabetes: the TODAY cohort at baseline. J. Clin. Endocrinol. Metab. 2011;96(1):159–167.

27. Dabelea D, Mayer-Davis EJ, Saydah S, Imperatore G, Linder B, Divers J, Bell R, Badaru A, Talton JW, Crume T, Liese AD, Merchant AT, Lawrence JM, Reynolds K, Dolan L, Liu LL, Hamman RF, SEARCH for Diabetes in Youth Study. Prevalence of type 1 and type 2 diabetes among children and adolescents from 2001 to 2009. JAMA 2014;311(17):1778–1786.

28. for Disease Control C, Prevention, Others. National diabetes statistics report, 2017. 2017. Available at: https://www.cdc.gov/diabetes/data/statistics/statistics-report.html.

29. Gast KB, Tjeerdema N, Stijnen T, Smit JWA, Dekkers OM. Insulin resistance and risk of incident cardiovascular events in adults without diabetes: meta-analysis. PLoS One 2012;7(12):e52036.

30. Krauss RM, Lindgren FT, Wingerd J, Bradley DD, Ramcharan S. Effects of estrogens and progestins on high density lipoproteins. Lipids 1979;14(1):113–118.

